# Aging-associated endolysosomal decline drives inflammaging and neurodegeneration through the STING-IFN-I axis

**DOI:** 10.64898/2026.03.15.711864

**Authors:** Maria Öberg, Ivana Maric, Anneli Strömberg, Caitlyn Myers, Najmeh Saffarzadeh, Daniela Fabrikova, Ivo Fabrik, Laia Rivas-Galvez, Karolina P Skibicka, Marzena Kurzawa-Akanbi, Gesine Paul, Nelson O. Gekara, Anetta Härtlova

## Abstract

All animals age. However, aging is a heterogeneous process and individual organisms age differently. Moreover, within the same organism, cells or organs do not age at the same time or speed. For instance, although neurodegeneration is a key trait of aging, neurological symptoms normally manifest long after multiple indicators of aging in peripheral tissues. The genetic determinants of aging remain poorly understood. Mutations in leucine-rich repeat kinase 2 (LRRK2) are major genetic risk factors for Parkinson’s disease (PD). By analyzing PD patients and mice with LRRK2 gain of function mutation (LRRK2^GoF^), we demonstrate that PD is an accelerated aging disease characterized by systemic low-grade STING-dependent inflammation (inflammaging) that first manifests in the periphery then disrupts the blood brain barrier and progresses to the brain resulting in neurodegeneration. Mechanistically, we demonstrate that a primary consequence of aging or *Lrrk2^GoF^* is endolysosomal decline. This results in the cytosolic build-up of extraneous self-DNA and subsequent shedding of DNA-containing extracellular vesicles thereby triggering the cGAS- STING pathway cell-intrinsically and intercellularly in distant host cells. This study unveils the cGAS-STING pathway and LRRK2^GoF^ as key determinants and potential targets for preventive or therapeutic strategies against accelerated aging, inflammaging and neurodegeneration.

## Introduction

Aging, a generalized and progressive decline in cellular and organ functions is an inevitable natural biological process that occurs in all animals^1,2^. However, aging is a heterogenous process and individual organisms age differently. Furthermore, within individual organisms, cells or organs do not age at the same time or speed^3–5^. For example, although neurodegeneration is a key trait of aging, neurological symptoms manifest long after several indicators of aging in peripheral tissues. In fact, even at advanced age, many people do not display overt cognitive impairment, suggesting a delay in age-associated decline in brain functions relative to peripheral organs.

An unavoidable outcome of aging is progressive low-grade inflammation (i.e., inflammaging). A growing body of evidence suggests that this smoldering inflammation is not only a consequence but also an accelerator of the aging process and related pathologies that affect different organs. In fact, although long considered an immune privileged organ, it is becoming increasingly apparent that age-associated decline in brain function could also be linked to inflammaging^6–15^. However, the molecular underpinnings of inflammaging and how this culminates in neurodegeneration and whether such inflammation starts and occurs exclusively in the brain, or starts elsewhere remain unresolved questions.

Parkinson disease (PD) is a progressive neurodegenerative disorder characterized by motor deficits, including resting tremor, bradykinesia, rigidity and postural instability^16^. Patients with PD also suffer from a variety of non-motor symptoms of which many affect peripheral organs (such as constipation). Those can even precede the onset of motor symptoms by decades and have been linked to an increased risk of developing PD^17–22^. While aging is the largest risk factors for developing PD, several genetic risk factors have also been identified. Key among these include mutations in the leucine-rich repeat kinase 2 (*LRRK2*) gene that encodes a large cytoplasmic protein (2527-amino acids). The most prevalent and best studied is the G2019S mutation that increases LRRK2 kinase activity^3,6–8,23^ and is associated with neuronal toxicity^23^. To date, the pathogenic mechanism behind LRRK2-driven PD progression and its link to aging remain unclear.

## Results

### Parkinson’s disease patients exhibit elevated age-associated systemic IFN-I

Spontaneous induction of type I interferons (IFN-Is) is a key hallmark of inflammaging^6–14^. In a blinded analysis of plasma samples from young (25-30 years) or aged (50-70 years old) healthy individuals or PD patients, we observed that aging-associated increase in systemic IFN-I and not NF-kB activity was a key feature distinguishing PD patients from healthy or young individuals (Fig. 1a-b, Supplementary Table 1). The highest IFN-I response was seen in a PD patient with a G2019S gain of function mutation in *LRRK2* (hereafter referred to as *LRRK2^GoF^*). We also found that compared to healthy individuals, blood monocytes from this patient had elevated transcripts for *IFNB1,* but not *TNFA* and that incubation with the LRRK2 small molecule inhibitor (MLI-2) could normalize the elevated *IFNB1* response to that in monocytes from healthy individuals (Fig. 1c-d). Consistent with the elevated IFN-I response, monocyte-derived macrophages from PD patients were more resistant to Herpes simplex virus (HSV-1) infection (Fig. 1d).

**Fig. 1:**
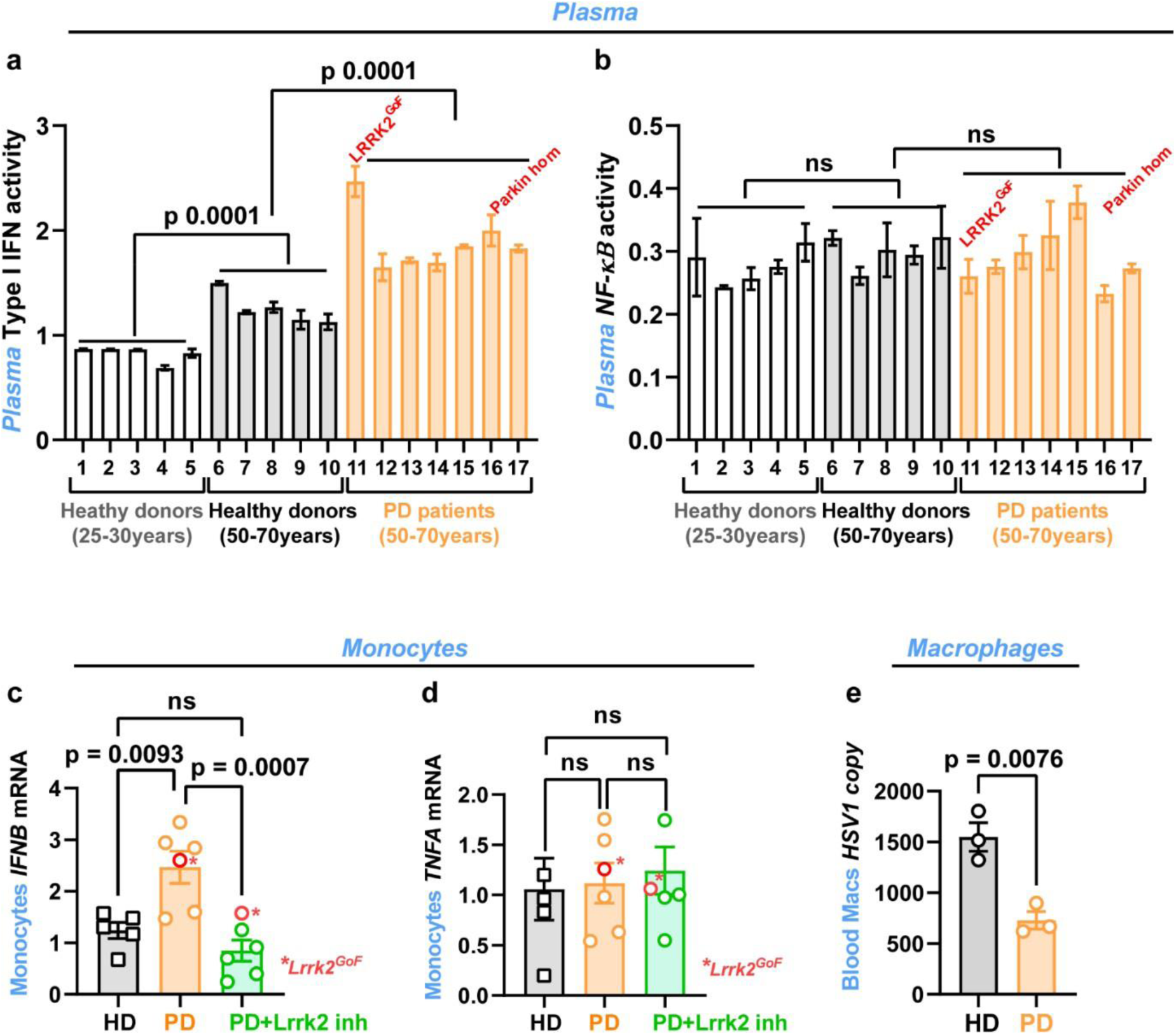
Parkinson’s patients exhibit elevated spontaneous systemic type I IFN. (**a, b**) Type I IFN (**a**) and NF-kB (**b**) activities in plasma from young (Age=25-30 years) and aged (Age=50-70years) healthy donors (HD) and PD patients (Age= 50-70; *Parkin homozygote mutation=24years), young/aged HD: n=5, PD: n=7. (**c-d**) Peripheral monocytes from sporadic PD patients and PD patient with PARKIN or LRRK2^GoF^mutation (Lrrk2 G2019S) exhibit elevated spontaneous type I IFN response. Transcripts of *IFNB1* **(c)** and *TNFA* (**d**) in monocytes from healthy donors (HD) or PD patients (PD) treated or not with the specific LRRK2 inhibitor MLI-2 for 6 hrs. HD: n=5, PD: n=6 (**e**) Monocytes-derived macrophages from PD patients *LRRK2^Go^*^F^ mutation are more resistant to herpes simplex virus (HSV1) infection. Data are shown as mean ± SEM of pooled HD and sporadic PD samples. HD: n=3, PD: n=3. Statistical tests: two-sided Student’s t-test (**e**) and one-way ANOVA followed by Tukey’s post hoc test (**a-d**). Error bars represent mean ± SEM.

### *Lrrk2^GoF^* accelerates aging and inflammaging in the peripheral tissues

In view of the above data, we next inquired whether *LRRK2^GoF^* also accelerates inflammaging in peripheral tissues of mice. Using a combination of IFN-I bioassay and qPCR, to analyze plasma, circulating monocytes, bone marrows, and spleens of 3 months vs. 12 months old wild-type and *Lrrk2^GoF^* mice, we observed that similar to humans, in mice, a hallmark of *Lrrk2^GoF^* was an increased age-associated IFN-I response (Fig. 2a-d). *Lrrk2^GoF^* mice also exhibited elevated inflammatory monocytes (CD11b^+^Ly6C^+^ high) in the blood (Fig. 2e, Extended Data Fig. 1) as well as the expression of the leukocyte activation marker CD86 (Fig. 2f). By bulk RNAseq we also observed that spleens of *Lrrk2^GoF^* mice had elevated expression not only of inflammatory but also senescence genes (Fig. 2g-h), prompting us to further inquire the impact of *Lrrk2^GoF^* on tissue senescence. Compared to wild-type, inguinal white adipose tissues (iWAT) from *Lrrk2^GoF^* mice had increased activity for the senescence- associated-beta-galactosidase (Fig. 2i-j). Together, these data demonstrate that a key consequence of *Lrrk2^GoF^* is accelerated aging and inflammaging in peripheral tissues.

**Fig 2:**
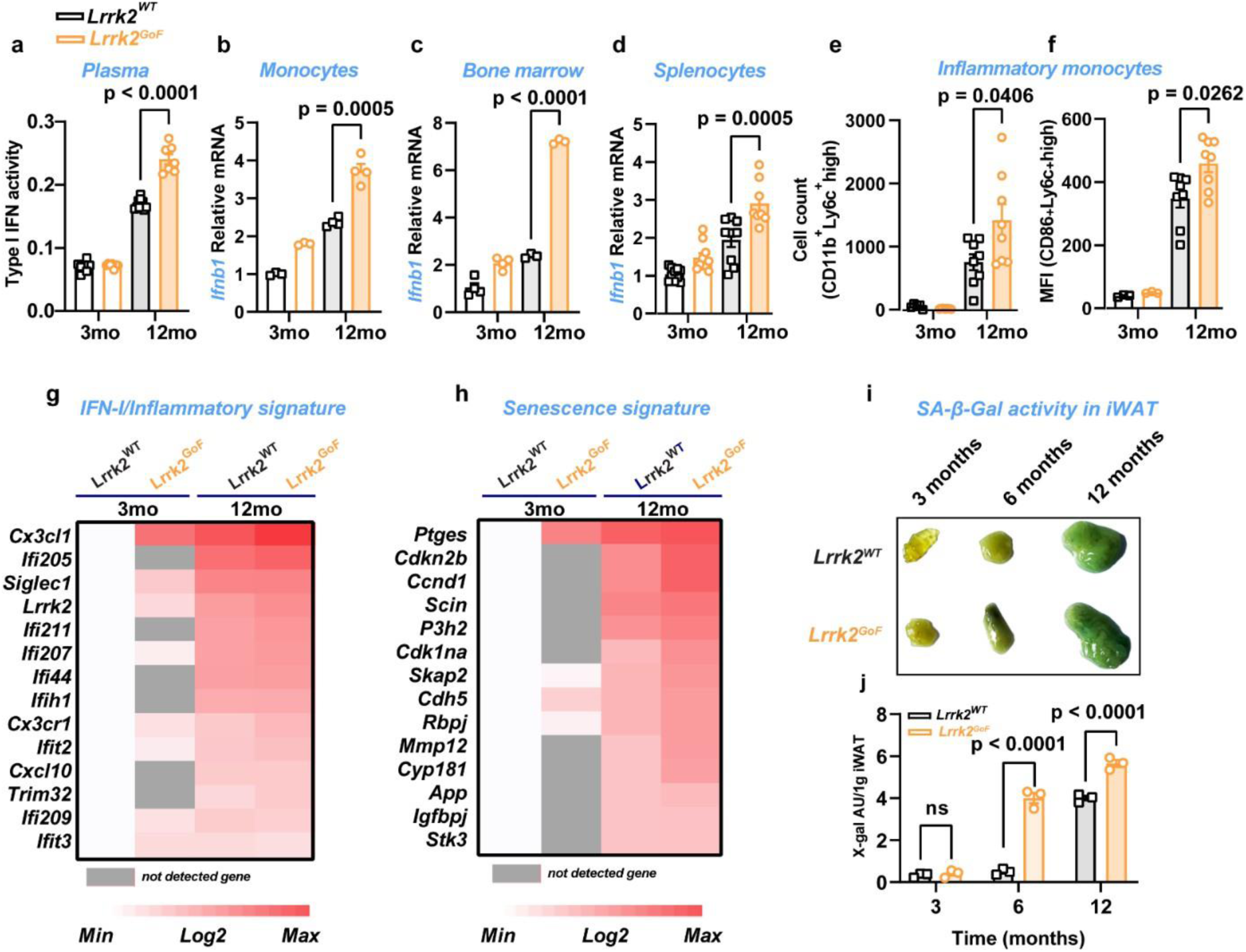
*Lrrk2^GoF^* mice exhibit accelerated age-associated chronic inflammation. (**a**) Type I IFN activity in plasma from 3 and 12 months old *Lrrk2^WT^* and *Lrrk2^GoF^* mice (n=7 mice per group). **(b-d)** *Ifnb1* transcripts in peripheral blood monocytes **(b)**, bone marrow **(c)** and splenocytes **(d)** of 3 and 12 months old *Lrrk2^WT^* and *Lrrk2^GoF^*mice. (**b, c**: n=3-4 mice per group) (**d**: n=8-12 mice per group) **(e)** Cell counts of CD11b^+^Ly6C^+^high inflammatory monocytes in blood of 3 and 12 months old *Lrrk2^WT^* and *Lrrk2^GoF^*mice. (n=8 mice per group). **(f)** Cell surface expression of the activation marker CD86 on the inflammatory monocytes in blood of 3 and 12 months old *Lrrk2^WT^* and *Lrrk2^GoF^* mice (n=8 mice per group). **(g-h)** Heatmap showing mean of differently expressed type I-IFN/inflammatory (**g**) and senescence (**h**) genes in splenocytes of 3 and 12 months old *Lrrk2^WT^* and *Lrrk2^GoF^* mice (n=3 mice per group).**(i)** SA-β-Gal activity in iWAT of *Lrrk2^WT^* and *Lrrk2^GoF^*mice. **(j)** Quantification of SA-β-Gal activity in iWAT of *Lrrk2^WT^* and *Lrrk2^GoF^*mice. (n=3 mice per group). Statistical tests: one-way ANOVA followed by Tukey’s post hoc test (**a-f, j**). Error bars represent mean ± SEM.

### *Lrrk2^GoF^* accelerates brain inflammaging and neurodegeneration

The brain is widely viewed as an immune privileged organ. However, growing body of evidence indicates that brain cells, for example microglia, are immune-competent and can mount an IFN-I response^24^. IFN-I signaling is essential for tissue homeostasis and neuroprotection^24,25^. However, if excessive, IFN-I signaling in the brain often results in neuropathology^14, 25–28^. Based on these results, we asked whether *Lrrk2^GoF^* also promotes spontaneous inflammation in the brain and whether this contributes to age-associated neurological decline. Using qPCR and bulk RNAseq, we found that compared to wild-type mice, microglia from *Lrrk2^GoF^* mice had elevated IFN-I signatures. This increase was particularly evident in the aged (12 months old) compared to the young (3 months old) mice, suggesting that *Lrrk2^GOF^* is in fact a risk factor for accelerated aging and associated brain inflammation (Extended Data Fig. 2a-b). Notably, when subjected to the Open Field and Rotarod tests, we observed that the age-dependent decrease in locomotor activity was exacerbated in *Lrrk2^GoF^* mice (Extended Data Fig. 2c-e). These data demonstrate that in addition to peripheral tissues, *Lrrk2^GoF^* is also a driver of aging-associated brain inflammation and neurodegeneration.

### *Lrrk2^GoF^*-driven inflammaging in the periphery precedes brain inflammaging and neurodegeneration

Aging affects all organs. However, the rate of aging and the associated decline in function likely varies from organ to organ^3,5^. As noted above, although neurodegeneration is a hallmark of aging, many individuals do not display overt neurological symptoms, even at advanced age. Of those who do, such symptoms normally become apparent long after the manifestations of several indicators of aging in peripheral tissues, suggesting that aging of the brain and peripheral tissues likely do not progress with the same kinetics. Inflammation is thought to be one of the major drivers of aging-related pathologies and decline in organ function^6–14,29^. Thus, to investigate a potential causal link between *Lrrk2^GoF^*-driven inflammaging and neurodegeneration, we compared the kinetics of peripheral versus brain inflammation and associated neurodegeneration. We found that although *Lrrk2^GoF^* mice already had a significantly higher IFN-I response in the periphery by the age of 3 months (Fig. 3a), this increase was not evident in the brain until the mice were 12 months old (Fig. 3b). Coincidentally, *Lrrk2^GoF^*-driven decline in locomotor coordination was also not apparent until 12 months (Fig. 3c). These data demonstrate that the onset of *LRRK2^GoF^*-driven inflammaging does not occur at the same rate in different organs, but first manifests in the peripheral tissues then progresses to the brain and therefore likely contributes to observed neuronal decline.

**Fig 3:**
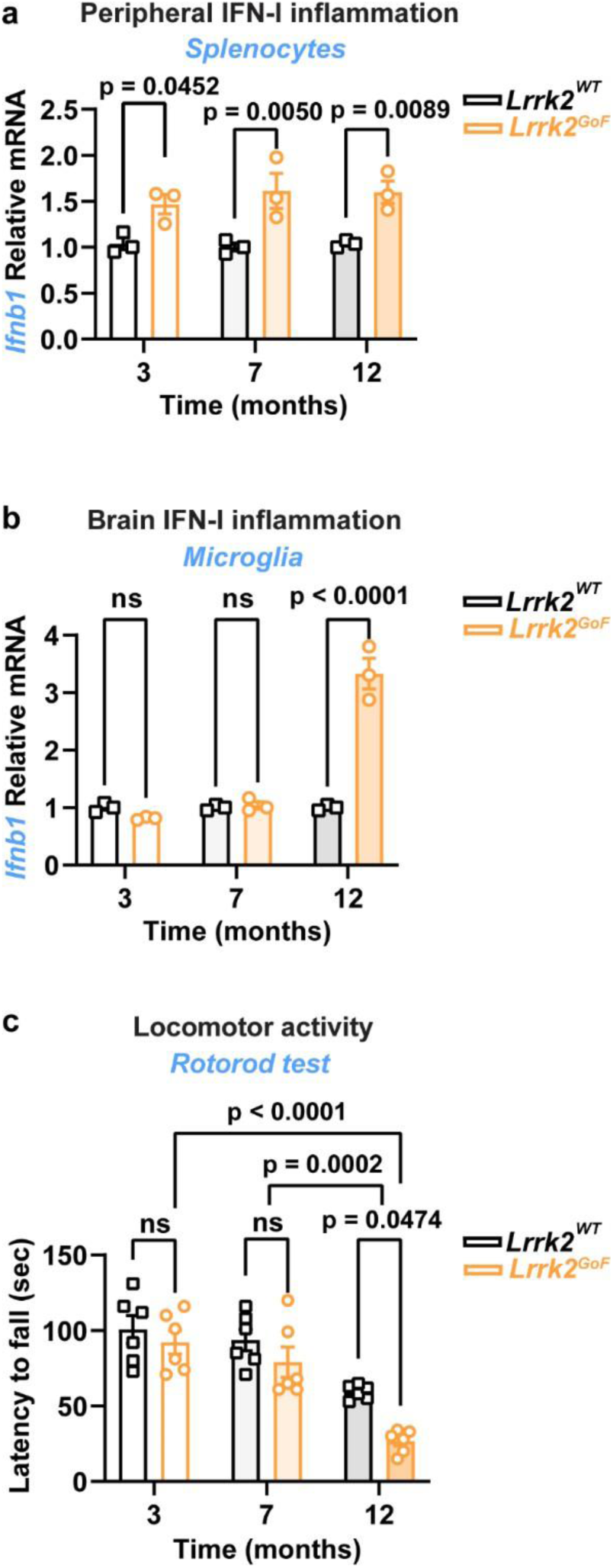
*Lrrk2^GoF^*-driven inflammaging in the peripheral tissues precedes brain inflammation and neurodegeneration. **(a-b)** *Ifnb1* transcripts in splenocytes **(a)** microglia **(b)** from 3, 7 and 12 months old *Lrrk2^WT^* and *Lrrk2^GoF^* mice. Data are show as comparison of *Lrrk2^GoF^* compared to wild-type counterparts for each time-point (n=3 mice per group). **(c)** Rotarod test showing locomotor activity in 3, 7 and 12 months old *Lrrk2^WT^* and *Lrrk2^GoF^*mice (n=6 mice per group). Statistical tests: one-way ANOVA followed by Tukey’s post hoc test **(a-c)**. Error bars represent mean ± SEM.

### *LRRK2^GoF^* accelerates inflammaging and neurodegeneration via STING signaling

Activation of the IFN-I response occurs via cell membrane-localized Toll-like receptors (TLRs) or intracellular innate immune receptors such as the Cyclic GMP-AMP synthase (cGAS)^29–33^. TLRs surveys the extracellular environment for different danger molecules and signal through key adaptors such as TRIF/TICAM (TIR-domain-containing adapter-inducing interferon-β)^33^. cGAS on the other hand senses DNA inside the cell and signals through the adaptor STING (Stimulator of interferon genes^34^. To elucidate the innate immune pathways involved in *Lrrk2^GoF^*-driven inflammaging and its potential link to neurodegeneration, we interbred *Lrrk2^GoF^* mice with those lacking TRIF or STING. We found that while the TLR-TRIF pathway had no effect, STING ablation completely reversed the elevated IFN-I response in *Lrrk2^GoF^* mice to that in wild-type controls (Fig. 4a, Extended Data Fig. 3). STING deficiency also reduced the expression of cellular markers of inflammation in microglia of aged mice including CD86 and MHCII (Fig. 4a-c). Importantly, when we subjected the mice to Open Field and Rotarod tests for locomotor activity, we found that STING ablation could protect *Lrrk2^GoF^* mice from age-associated neurological decline (Fig. 4d-f), demonstrating that *Lrrk2^GoF^*-driven inflammaging and neurodegeneration were due to spontaneous activation of the cGAS-STING pathway.

**Fig. 4:**
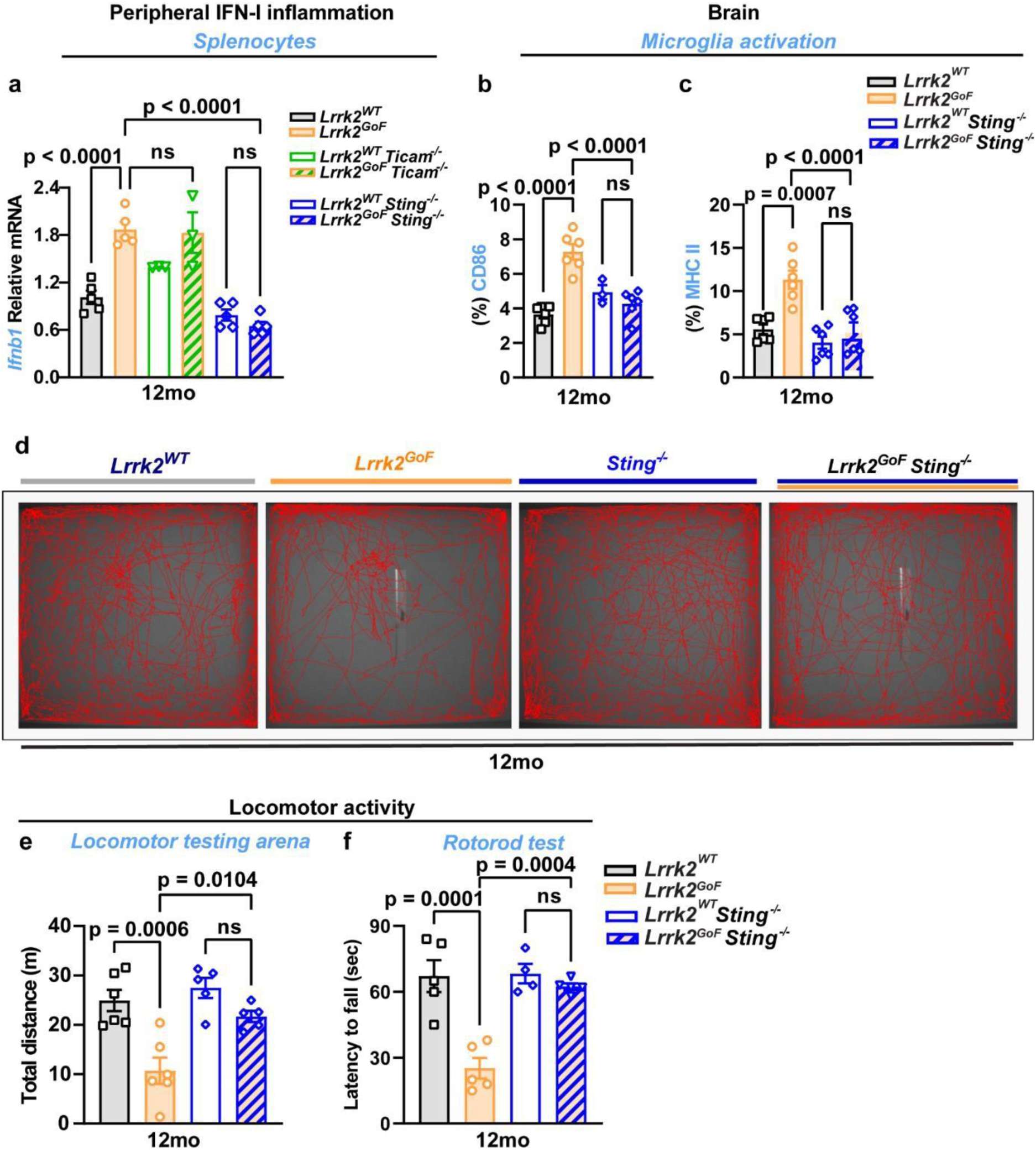
*Lrrk2^GoF^* accelerates age-associated neurological decline via the Sting pathway. (**a**) *Ifnb1* transcripts in splenocytes of 12 months old *Lrrk2^WT^* and *Lrrk2^GoF^*mice lacking or not *Ticam* or *Sting* signaling pathway (n=3-5 mice per group). **(b-c)** Cell surface expression of the activation marker CD86 **(b**) and MHC-II **(c)** on microglia of 12 months old *Lrrk2^WT^* and *Lrrk2^GoF^*mice lacking or not *Sting* signaling pathway (n=3-6 mice per group). **(d)** Representative images of track plot reports recorded during the test sessions (ANY-maze) of 12 months old *Lrrk2^WT^* and *Lrrk2^GoF^*mice lacking or not *Sting* signaling pathway (n=5-6 mice per group). **(e)** Locomotor activity of 12 months old *Lrrk2^WT^* and *Lrrk2^GoF^*mice lacking or not *Sting* signaling pathway measured as total distance travelled within open field area (n=5-6 mice per group). **(f)** Rotarod test showing locomotor activity of 12 months old *Lrrk2^WT^* and *LRRK2^GoF^*mice lacking or not *Sting* signaling pathway (n=4-5 mice per group). Statistical tests: one-way ANOVA followed by Tukey’s post hoc test **(a-c, e, f)**. Error bars represent mean ± SEM.

### *Lrrk2^GoF^* accelerates senescence and release of inflammatory extracellular vesicles

In view of the above demonstration that *Lrrk2^GoF^* is associated with accelerated inflammaging and senescence *in vivo* (Fig. 2), next we sought to directly assess the effect of *Lrrk2^GoF^* on senescence in cell culture. Upon culture for several weeks, we found that mouse embryonic fibroblasts (MEFs) from *Lrrk2^GoF^* mice stopped proliferating earlier than the wild type MEFs (*Lrrk2^WT^*) (Fig. 5a) and exhibited a higher expression of the senescence marker beta-galactosidase (Fig. 5b), and an enhanced *Ifnb1* that was reversible in the presence of the Lrrk2 inhibitor MLI-2 (Fig. 5c).

**Fig 5:**
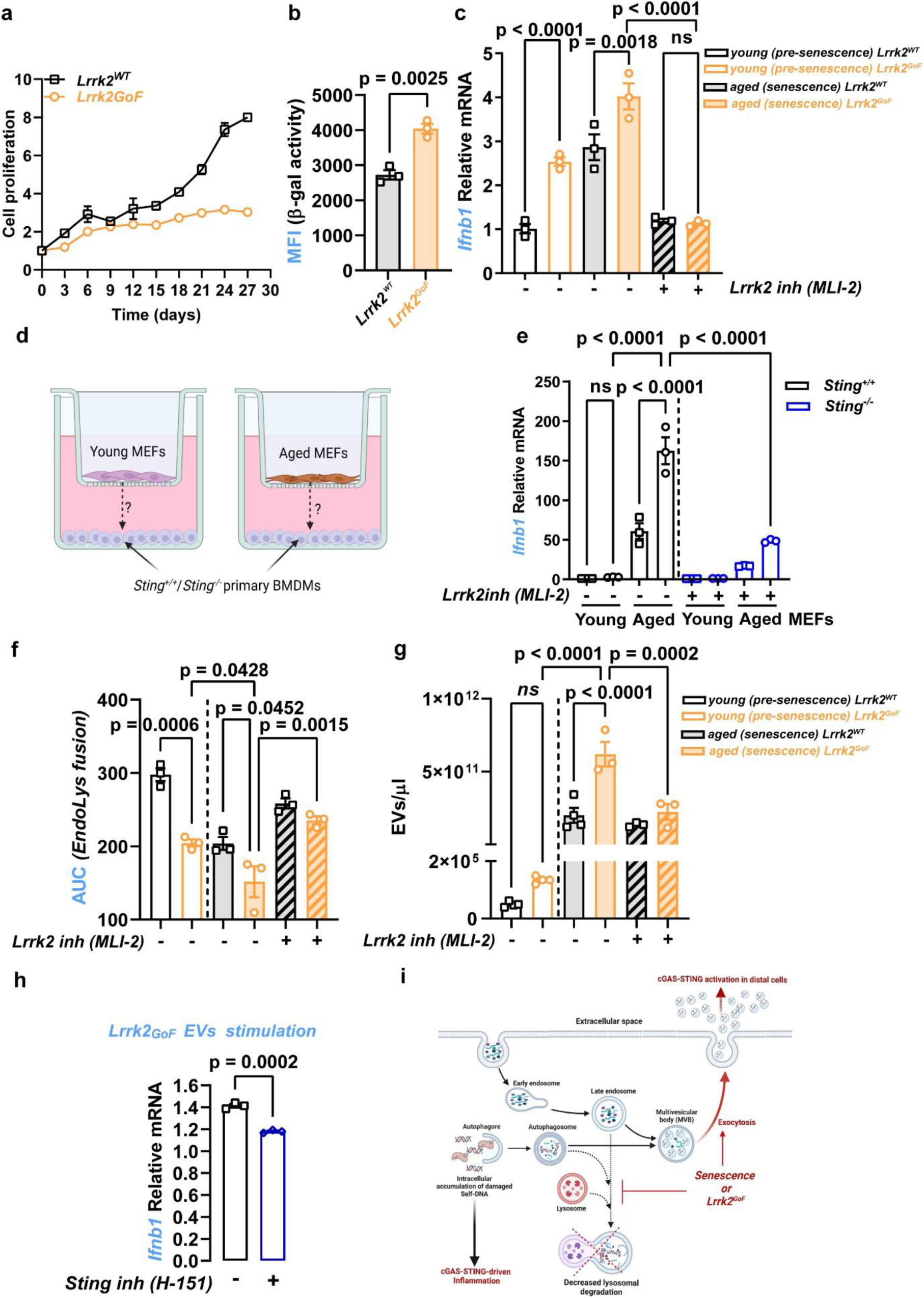
*Lrrk2^GoF^* accelerates senescence and release of inflammatory extracellular vesicles. (**a**) Proliferation curve of primary *Lrrk2^WT^* and *Lrrk2^GoF^* mouse embryonic fibroblasts (MEFs). **(b)** The mean fluorescence intensity of SA-β-Gal-positive aged *Lrrk2^WT^* and *Lrrk2^GoF^* MEFs (n=3 per group). **(c)** *Ifnb1* transcripts in young (pre-senescent) and aged (senescent) *Lrrk2^WT^* and *Lrrk2^GoF^* MEFs treated or not with the Lrrk2 specific inhibitor MLI-2 for 24h (n=3 per group). **(d)** Schematic depicting trans-well co-cultures of young and aged *Lrrk2^WT^* and *Lrrk2^GoF^* MEFs with either *Sting^+/+^* or *Sting^-/-^* bone marrow-derived macrophages (BMDMs). **(e)** *Ifnb1* transcripts in recipient *Sting^+/+^* and *Sting^-/-^* BMDMs co-cultured with young or aged *Lrrk2^WT^* and *Lrrk2^GoF^* in trans-wells for 24h (n=3 per group). **(f)** Endo-lysosomal BSA DQ-green degradation in young and aged *Lrrk2^WT^* and *Lrrk2^GoF^* MEFs pre-treated or not with 100nM Lrrk2 inhibitor MLI-2 for 1h, AUC: area under the curve (n=3 per group). **(g)** Quantification of extracellular vesicles release of young and aged *Lrrk2^WT^* and *Lrrk2^GoF^* MEFs treated or not with Lrrk2 inhibitor MLI-2 for 48h by nano-particle tracking analysis (NTA) (n=4 per group). **(h)** *Ifnb1* transcripts in young *Lrrk2^W^*^T^ MEFs with aged *Lrrk2^GoF^*-derived extracellular vesicles for 8h pre-treated or not with Sting inhibitor H-151 (n=3 per group). **(i)** Schematic illustration of how inhibition of endolysosomal clearance by *LRRK2^GoF^* results in spontaneous activation of cGAS-STING-IFN-I signaling cell intrinsically and in distal cells. Statistical tests: two-sided Student’s t-test **(b, h)**, one-way ANOVA followed by Tukey’s post hoc test **(a, c, e-g)**. Error bars represent mean ± SEM.

In the course of these analyzes, we observed that when co-cultured with primary bone marrow derived macrophages (BMDMs) in trans-wells (to prevent direct cell - cell interactions but to allow indirect communication via diffusion of nanometer sized particles), senescent MEFs (WT or *Lrrk2^GoF^*) were able to evoke a STING-dependent IFN-I response in distant BMDMs (Fig. 5d-e). Together, these data demonstrated 1) that *Lrrk2^GoF^*-driven inflammation is due to accelerated senescence and 2) that aging or *Lrrk2^GoF^*-driven STING- IFN-I signaling is not only a cell-intrinsic event but that *Lrrk2^GoF^* or senescent cells can also activate the same pathway in distal cells remotely.

The progressive accumulation of damaged self-DNA in the cytosol as a result of genome instability^35^ or mitochondrial dysfunction^36^ is a key trait of senescence. Inflammaging is widely assumed to be due to the cell-intrinsic cGAS-STING sensing of such DNA within senescent cells^35, 37–39^. However, our result show that cGAS-STING signaling is not restricted to senescent cells but spreads to distal bystander cells. What is the molecular basis for this remote activation of cGAS-STING?

Extracellular vesicles (EVs) are nanosized lipid bilayer enclosed vesicles (30- 100 nm) of endocytic origin secreted by most cell types and contain different biomolecules, including DNA^40^. Previously, we showed that *Lrrk2^GoF^* cells are defective in endolysosomal clearance^41^. This together with previous reports suggesting an inverse correlation between endolysosomal clearance and EVs release^42–45^, prompted us to inquire whether *Lrrk2^GoF^*-driven inflammaging was related to an imbalance in endolysosomal clearance versus EVs release.

We reasoned that by impeding endolysosomal clearance, LRRK2^GoF^ not only results in increased accumulation of damaged self-DNA intracellularly, but also in an increased release of DNA-containing EVs, which in turn could deliver DNA into distal recipient cells, and that both of these characteristics were likely responsible for the cell-intrinsic and remote activation of the STING-IFN-I axis in distal cells, respectively. In line with this hypothesis, we found that compared to wild-type MEFs, *Lrrk2^GoF^* MEFs were defective in endolysosomal activity (Fig. 5f, Extended Data Fig. 4a) and released more EVs in an age-dependent manner (Fig. 5g, Extended Data Fig. 4b). Incubation with the LRRK2 inhibitor MLI-2 restored endolysosomal activity and EVs release in *Lrrk2^GoF^* to that in *Lrrk2^WT^* cells (Fig. 5f-g). These EVs contained DNA and could transfer it to primary BMDM, thus triggering an IFN-I response in a STING dependent manner (Fig. 5h, Extended Data Fig. 5a, b).

Viewed alongside the above data demonstrating that *Lrrk2^GoF^* is an accelerator of senescence (Fig. 5a-b), we asked whether senescence also influences endolysosomal clearance independently of the *Lrrk2* mutation. By comparing young and aged *Lrrk2^WT^* MEFs, we found that endolysosomal activity was decreased in senescent cells (Extended Data Figure 5c), implying that progressive decline in endolysosomal clearance is a key trait of cellular aging, an idea is consistent with previous observations^46–49^. Taken together, these results demonstrate that *Lrrk2^GoF^* is a driver of senescence and that defective endolysosomal clearance and increased release of DNA-containing EVs are consequences of the aging process. In addition, we show that age-driven cGAS-STING-IFN-I signaling is not only restricted within senescent cells, but that by shedding DNA-containing EVs, senescent cells can remotely activate the cGAS-STING-IFN-I axis in distal cells, thus suggesting how senescent cells could propagate inflammation within and across tissues (schematically summarized in Fig. 5i).

### Aging and *LRRK2^GoF^*accelerates endolysosomal decline and release of inflammatory EVs in mice and humans

To further elucidate the mechanisms underlying LRRK2^GoF^-driven inflammaging, we inquired whether *Lrrk2^GoF^*also affects endolysosomal clearance and promotes EVs release *in vivo*. When we compared primary cells from *Lrrk2^WT^* and *Lrrk2^GoF^* mice, we found that blood monocyte-derived macrophages from *Lrrk2^GoF^* mice had a decreased endolysosomal proteolysis rate (Fig. 6 a-b). Consistently, *Lrrk2^GoF^* mice also exhibited an increased age-dependent accumulation of DNA-containing EVs in the blood and the cerebrospinal fluid (Fig. 6c-d, Extended Data Fig. 7a-d). We noted that, while an increase in EVs accumulation in the blood of *Lrrk2^GoF^* mice was already evident in young mice (3 months old), accumulation in the cerebrospinal fluids was not apparent until mice were older (12 months old) (Fig. 6c-d). Interestingly, this delayed accumulation of EVs in cerebrospinal fluids coincided with the elevated IFN-I response and expression of cellular markers of inflammation in the brain and the onset of neurological symptoms (Extended Data Fig. 2, Fig. 3, Fig. 5). Upon incubation with BMDM, EVs isolated from aged *Lrrk2^GoF^* mice induced an IFN-I response in a STING dependent manner (Fig. 6e-f), suggesting a causal link between EVs accumulation and *Lrrk2^GoF^*-driven cGAS-STING-dependent inflammaging.

**Figure 6:**
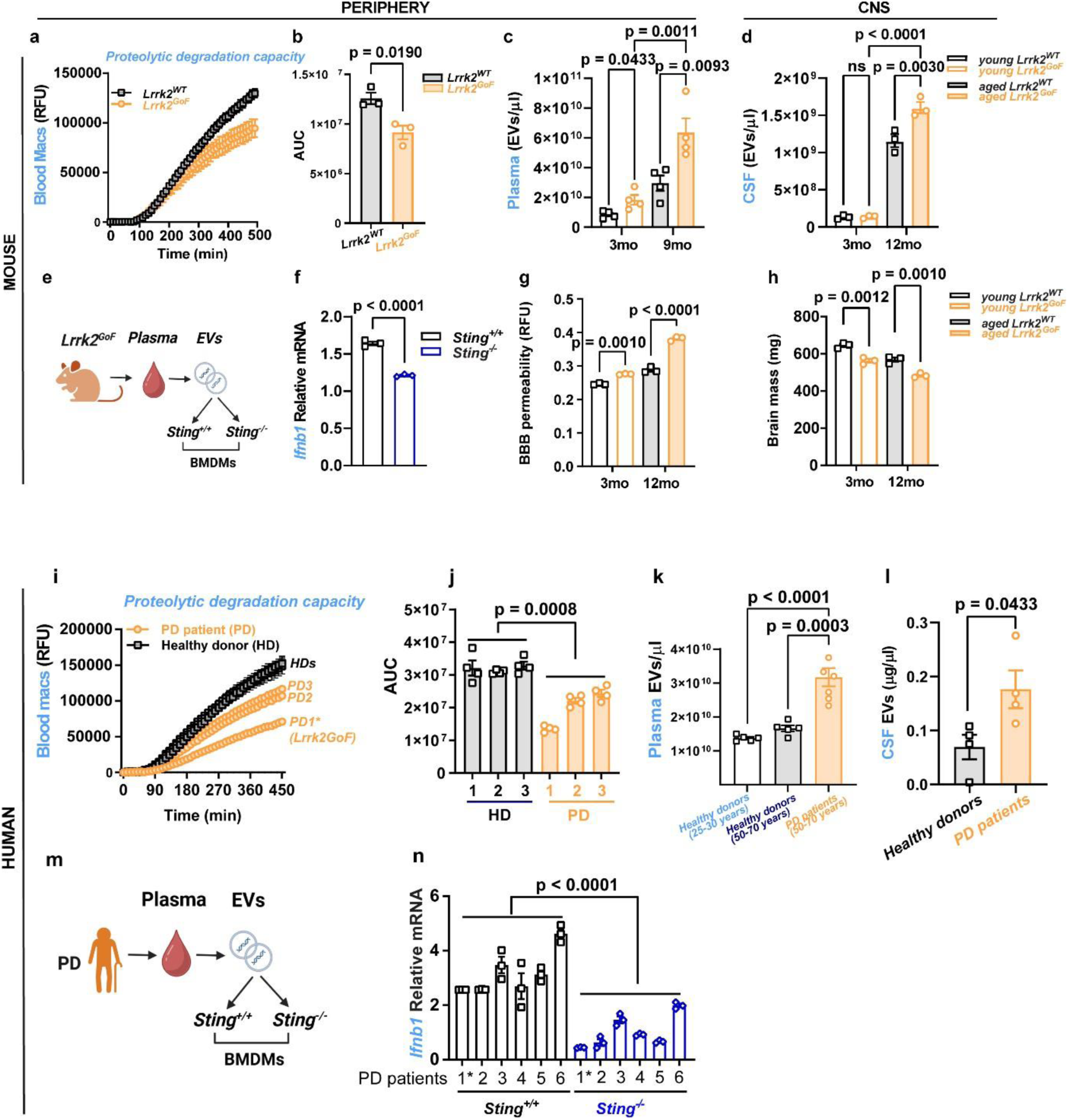
L*R*RK2GoF accelerates age-associated release of DNA-containing extracellular vesicles in mice and humans. (**a**) Endo-lysosomal BSA DQ-green degradation in aged *Lrrk2^WT^* and *Lrrk2^GoF^* blood monocyte-derived macrophages (n=3 mice per group). **(b)** Area under the curve (AUC) of the endo-lysosomal proteolytic activity in aged *Lrrk2^WT^* and *Lrrk2^GoF^* blood monocyte-derived macrophages (n=3 mice per group). **(c-d)** Quantification of extracellular vesicles (EVs) release in plasma **(c)** and cerebrospinal fluid **(d)** of young and aged *Lrrk2^WT^* and *Lrrk2^GoF^* mice by nanoparticle tracking analysis (NTA). **(e)** Schematic outline of stimulating *Sting^+/+^* or *Sting^-/-^* BMDMs with *Lrrk2^GoF^* EVs. **(f)** *Ifnb1* transcripts in *Sting^+/+^* or *Sting^-/-^* upon mouse *Lrrk2^GoF^* EVs stimulation for 6h (Isolated EVs from n=6-8 mice per group, pooled in 3 replicates). **(g)** Estimation of blood-brain barrier (BBB) permeability by Evans blue staining in young and aged *Lrrk2^WT^* and *Lrrk2^GoF^* brains (Isolated EVs from n=6-8 mice per group, pooled in 3 replicates). **(h)** Brain mass of young and aged *Lrrk2^WT^* and *Lrrk2^GoF^* brains (n=3 mice per group). **(i)** Endo-lysosomal BSA DQ-green degradation in blood monocyte-derived macrophages of healthy donors (HD) and Parkinson’s patients (PD) (n=3 donors per group). **(j)** Area under the curve (AUC) of the endo-lysosomal proteolytic activity in blood monocyte-derived macrophages of HD and PD (n=3 donors per group). **(k-l)** Quantification of EVs release in plasma **(k)** and cerebrospinal fluid **(l)** of HD and PD determined by either nanoparticle tracking analysis (NTA) or protein concentration (*Plasma*: n=5 HDs, 6 PDs; *CSF*: n=3 HDs, 3 PDs). **(m)** Schematic outline of stimulating *Sting^+/+^* or *Sting^-/-^* BMDMs with PD patients-derived EVs **(n)** *Ifnb1* transcripts in *Sting^+/+^* or *Sting^-/-^* in response to PD patients-derived EVs stimulation for 6h (*Plasma-enriched EVs*: 6 PDs). Statistical tests: Student’s t-test **(b, h, l)**, one-way ANOVA followed by Tukey’s post hoc test **(c-e, f, j, k, n**). Error bars represent mean ± SEM.

The blood-brain barrier (BBB) that limits the unregulated exchange of macromolecules and cells between the blood and the central nervous system is essential for maintaining the immune privileged status of the brain. Acute peripheral inflammation, for example due to acute infection can disrupt the integrity of the BBB^50–55^. Therefore, we inquired whether the progressive low-grade inflammaging in the periphery, also affects BBB integrity. For this, we injected Evans blue dye intravenously and monitored its accumulation into the brain. We found that aging and *Lrrk2^GoF^* were indeed associated with increased BBB permeability and decreased brain mass (Fig. 6g-h). Together, these results suggest that increased inflammation and DNA-containing EVs in the periphery could spread to the CNS and contribute to brain inflammation.

Finally, to verify these findings in clinically relevant settings, we compared samples from healthy individuals with those from PD patients. Similar to observations in mice (Fig. 6a-b), blood monocytes from PD patients (with unknown or known genetic risk factors including PARKIN^56^ and *LRRK2^GoF^*) showed a decrease in endolysosomal degradation (Fig. 6i-j). Furthermore, we found that in humans, aging was associated with increased accumulation of EVs in the blood and cerebrospinal fluids and that this was more pronounced in PD patients (Fig. 6k-l, Extended Data Fig. 7e). When co-incubated with BMDM, EVs isolated from PD patients could induce an IFN-I response in a STING-dependent manner (Fig. 6m-n). In summary, these data reveal *LRRK2^GoF^* as an accelerator of aging and inflammaging. Moreover, these data establish enhanced release of DNA-containing inflammatory EVs as a key hallmark of aging, thus suggesting how senescence-associated inflammation could spread within and across tissues, including from the periphery to the brain.

## Discussion

The genetic determinants of aging are still poorly understood. *LRRK2^GoF^* mutations are a major risk factor for Parkinson’s disease (PD). By analyzing PD patients and *Lrrk2^GoF^* genetic mice, our data suggest that PD is an accelerated aging disease characterized by systemic low-grade inflammation that first manifests in the periphery, then with age disrupts the blood-brain barrier and progresses to the CNS, ultimately resulting in neurodegeneration. Our data demonstrate that *Lrrk2^GoF^*-driven acceleration of aging, inflammaging and neurodegeneration involves activation of the STING-IFN-I axis. Furthermore, we demonstrate that a consequence of aging or *Lrrk2^GoF^* is a decrease in endolysosomal function. Our data support a model in which by impeding endolysosomal clearance, aging or *Lrrk2^GoF^* results not only in the intracellular accumulation of cellular waste including self-DNA culminating in cell-intrinsic activation of the STING pathway, but also in the release of DNA-containing EVs that mediate remote activation of the same pathway in distal host cells (schematic illustration Fig. 5i).

Although most PD cases are often due to unknown risk factors^57^, we found that in addition to those with mutation in *LRRK2,* all the other PD patient in our cohorts also exhibit diminished endolysosomal clearance and an increased release of DNA-containing EVs into the blood and cerebrospinal fluids (Fig. 6i-l). Many genes associated with increased risk of PD such as PARKIN^58^, PINK1^59^, VPS35^60,61^, GBA^62^, SYT11^63,64^, ATP13A2/PARK9^65–67^, TMEM175^68,69^ have also been linked to endolysosomal dysfunction. These together with the present results suggest that diminished endolysosomal clearance, increased EVs release and associated activation of the STING-IFN-I axis might be common denominators not just for PD but aging in general.

PD is largely viewed as a progressive neurodegenerative movement disorder. However, PD patients have long been observed to display various inflammatory phenotypes and non-motor symptoms that can precede the disease diagnosis by decades^17–22^. In fact, inflammatory diseases involving peripheral organs are considered as major risk factors for developing PD. On the other hand, genetic variants such as LRRK2^GoF^ traditionally associated with PD are also risk factors for inflammatory disorders, for example inflammatory bowel disease^70–73^. These together with the present findings reinforce the idea that PD is not simply a brain-specific disease, but a systemic inflammatory disorder that might start in peripheral organs and then with age progresses to the brain. In other words, while the brain is immune privileged, analogous to a safe within a burning house, in the midst of smoldering inflammation in periphery tissues, with age, the brain cannot remain safe. Our demonstration that a key outcome of senescence is increased accumulation of DNA in the cytosol, as well as its release and delivery to distal host cells via EVs illustrates how age-driven inflammation is initiated within senescent cells and how it can spread within and across tissues, including from the periphery to the brain.

As noted above, PD is often associated with spectrum of comorbidities that precede the onset of motor symptoms and which may increase the risk of PD. However, to date the management strategies for PD have predominantly focused on restoring the dopaminergic system to treat motor symptoms and less on how to prevent or alleviate non-motor related comorbidities. Therefore, understanding the underlying mechanisms of PD and identifying predictive biomarkers at the premotor stages of the disease hold future promise for timely stratification of at-risk individuals for early interventions so as to prevent or delay disease progression. The demonstration herein that PD is an age-driven systemic inflammatory disease that starts in the periphery then progresses to the central nervous system (CNS) and that this involves the STING-IFN-I axis, and that LRRK2 is an accelerator of aging and associated inflammation represents a significant advance in our molecular understanding of the aging process and associated neurodegeneration. Additionally, it reveals the STING pathway and LRRK2^GoF^ as potential biomarkers and targets for preventive or therapeutic strategies.

## Supporting information

Resource Table

## METHODS

### Clinical samples ethics statement

Experiments involving human subjects were done according to the recommendations of the ethics committee as approved in permit Dnr2011/89. Informed consent was obtained from PD and healthy patients in compliance with the Helsinki declaration act.

### Mice ethics statement

Mice were maintained under specific pathogen free conditions and all experiments were approved and carried out according to the guidelines set by the ethical committee Vetenskapsrådet approval 2947/20.

### Mice

*Lrrk2^WT^* (*C57BL/6*) mice and *Lrrk2^GoF^* harbouring G2019S mutation (*C57BL/6-Lrrk2tm4.1Arte*) mice were purchased from Taconic (US). *Sting^-/-^* (*C57BL/6J-Tmem173gt/J*) and *Ticam^-/-^*(*C57BL/6J-Ticam1Lps2/J*) mice were purchased from the Jackson Laboratory. Mice were bred and crossed with each other at Gothenburg University to generate the following mouse lines *Lrrk2^W^Sting^-/- T^ or Lrrk2^GoF^Sting^-/-^ and Lrrk2^WT^Ticam^-/-^ or Lrrk2^GoF^ Ticam^-/-^*.

Adult mice were housed in specific pathogen free conditions with access to food and water ad libitum were used in all experiments. All animal experiments were conducted during the light-phase. All animal experiments were performed at Gothenburg University animal facility according to the guidelines set by the ethical committee (Vetenskapsrådet, approval no. 2947/20).

### Mouse embryonic fibroblasts (MEFs)

Primary MEFs were prepared from a pregnant *Lrrk2^WT^* and *Lrrk2^GoF^* mice at 13 or 14 days post-coitum. MEFs after passage 2 (P2) were collected and maintained as stock cells. MEFs were maintained in Dulbecco’s modified Eagle’s medium (DMEM, Gibco) supplemented with 10% fetal bovine serum (FBS), 2 mM l-glutamine, 1% penicillin-streptomycin.

### Interferon cell-based assay

**B16-Blue^TM^ ISG cells** (Invivogen, #bb-ifnabg), B16 F1melanoma cells, were maintained in DMEM with 10% heat inactivated FBS, 1% pen-strep, 100µg/ml Normocin and 100µg/ml Zeocin. **HEK-Blue^TM^ ISG cells** (Invivogen, #hkb-isg-1), human embryonic kidney cell line, were maintained in DMEM with 10% heat inactivated FBS, 1% pen-strep, 100µg/ml Normocin and 100µg/ml Zeocin. Both cell lines express the secreted embryonic alkaline phosphatase (SEAP) reporter gene under the control of the I-ISG54 promoter which is comprised of the IFN-inducible ISG54 promoter enhanced by a multimeric ISRE. Stimulation of B16- Blue™ ISG cells with IFNs, CDNs, such as cGAMP, or type I IFN inducers, such as transfected poly(dA:dT), triggers the activation of the I-ISG54 promoter and the production of SEAP.

### Bone marrow derived macrophages (BMDMs)

BMDMs were generated by culturing the mouse bone marrow cells in IMDM medium (GIBCO, Life Technologies) supplemented with 10% FCS (GIBCO, Life Technologies), 100 U/ml penicillin (Sigma-Aldrich), and 100 g/ml streptomycin (Sigma-Aldrich), 2 mM glutamine (Sigma-Aldrich) and 15% (v/v) L929 conditional medium.

### Primary microglia isolation

Mouse primary microglia cells were isolated by dissociating brain tissue using the Milteny Adult Brain dissociation kit (Milteny biotech, #130-107-677). Followed by isolation of brain microglia cells via magnetic isolation using CD11b positive Microbeads (Milteny biotech, #130-049-601).

### Human monocyte isolation

Peripheral Blood Mononuclear cells (PBMCs) were isolated from healthy and PD patient blood using Ficoll density gradient centrifugation at 800xg for 15min at RT. The PBMC were collected and resuspended in PBS/2mM EDTA then pelleted by centrifugation at 300xg for 8min at RT to eliminate platelets. Erythrocytes were removed by addition of 1x RBC lysis buffer. PBMCs were washed in PBS and monocytes were isolated by the use of CD14 positive magnetic beads (Milteny, #130-050-201).

### Viruses and viral propagation

Herpes simplex virus type I strain KOS KOS/Dlux/OriL (HSV-I-luc) containing a firefly luciferase reporter (Summers and Leib, 2002) was received from Nelson Gekara and propagated as previously described [74].

### Behavioral studies

In the **open field locomotor activity test**, mice were placed in an open box (35 x 35 cm) where total activity, discrete movements and distance traveled were measured. Total distance travelled indicates general capacity of the mice to locomote. Mice were placed in the center of the arena and allowed to explore freely during 25 minutes. Distance travelled, number of entries in center and time spent in center was scored using ANY-Maze video-tracking [75].

The **rotarod** test evaluates the motor performance of mice by measuring their ability to stay on a rotating cylinder (rotometer from Harvard Apparatus Panlab). Mice were trained to remain on the cylinder at the lowest speed (4 rpm) for one minute before qualifying for the actual test. In the test session, the cylinder accelerated from 4 rpm to 40 rpm over 5 minutes. The recorded data included the time and speed at which the mouse fell off.

### Blood Brain Barrier permeability

The blood brain barrier integrity was studied by Evans-blue. 2% Evans-blue (3ml/kg) was injected intravenous into the tail vein of three 3- and 12-month-old WT and *Lrrk2^GoF^* mice. After 1h Intracardiac perfusion was performed through the left ventricle with 20ml of 0.9% saline to remove intravascular Evans blue. Blood was collected from the right ventricle by cardiac puncture. Brain was dissected, weighed, and homogenized in 500µl 50% trichloroacetic acid. The samples were centrifuged at 10000xg at 4°C for 10 min. The supernatant was collected, and vacuum concentrated to 100µl. Fluorescence was measured at excitation 620nm and emission 680nm and normalized to the tissue weight and expressed as RFU/100mg brain tissue.

### Collection of Cerebrospinal fluid

Cerebrospinal fluid (CSF) was collected from cisterna magna during the terminal experiment. Animals were anesthetized with an intraperitoneal (i.p.) ketamine (90 mg/kg) and xylazine (10 mg/kg) injection. The collection was performed as shown in Lim *et al* [76]: a sharpened glass capillary was set up in a micromanipulator and attached to a thin tube connected to a syringe with a three-way valve. The anesthetized mouse was fixed in a stereotaxic apparatus with nose pointing down, positioning the back of the neck in a flat plane. Using scissors and forceps, the skin and muscle of the neck were cut to expose the cisterna magna at the base of the skull. The dura of cisterna magna was punctured with the capillary tube, and the setup was left in this position until CSF stopped drawing out or blood contamination was observed. Following CSF collection mice were directly terminated by cervical dislocation.

### Flow cytometry of circulating leukocyte subsets in mouse

Total number of myeloid cells, monocyte subsets and neutrophils were identified as depicted in Figure S1B. Blood was collected via cardiac puncture into ethylene-diaminetetraacetate (EDTA) lined tubes and immediately placed on ice. Red blood cells (RBCs) were lysed (3 times for 5 minutes) and leukocytes were centrifuged, washed and re-suspended in flow cytometry buffer for antibody staining (PBS containing 1% bovine serum albumine (BSA), 5mM EDTA and 0.05% NaN_3_). Samples were stained with FACS antibody for 1h at 4°C (antibody panels are in Table 1) in the dark then washed in PBS and resuspended in FACS buffer. Cells were analyzed on the LSRFortessa X-20 (BD) and FlowJo software, v 10.8.1.

| **Flow cytometry panels** | **Tissue** | **Antibodies** |
| --- | --- | --- |
| **Panel 1** | Brain, microglia | CXCR1-FITC, Clec12A-PE, CD11b-PE-cy7, CD206-AF647, CD45-BUV421, CD11c-BV711. |
| **Panel 2** | Brain, activated macrophages | MHCII-AF700, CD86-PE, Ly6C-BV605, CD45-BV421. |
| **Panel 3** | Blood | Brilliant stain buffer, TCRβ-PE-CF594, CD4-BUV421, CD8-BV711, B220-BUV496, CD11c-APC-Cy7, CD11b-BV510, CD45-FITC, 7AAD, F4/80-BUV395, Gr1-BV650, MHCII-AF700, CD86-PE, Ly6c-BV605, CD44-BV786, CD62L-APC, CD69-BUV737, Fc-block |

### SA-β-Gal staining in iWAT

Whole mount SA-β-Gal stainings on fat were performed using a kit (#9860, Cell Signaling), with a 20-min fixation followed by immersing samples in the staining solution at 37 °C for 12 h (iWAT). Quantification of X-gal activity was measured by absorbance at 615nm and normalized on 1g of a tissue.

### Mouse serum and CSF Extracellular vesicle (EVs) isolation

Mouse serum extracellular vesicles were isolated using the miCURY serum/plasma exosome isolation kit (Qiagen, #76603) according to manufacturer instructions. Mouse CSF extracellular vesicles were isolated using the miCURY cell/CSF/urine exosome isolation kit (Qiagen, #76743) according to manufacturer instructions.

### Human serum, CSF and MEFs supernatant extracellular vesicle (EVs) isolation

Extracellular vesicles from human serum, CSF and MEFs were isolated as described in Kurzawa-Akanbi et al [77]. Briefly, MEFs for extracellular vesicles isolation were grown in DMEM with 10% EVs depleted FBS (Thermo Fisher Scientific), 1% pen-strep. Serum from human and cell supernatant from MEFs were concentrated for extracellular vesicle isolation using 10 or 30kDa Amicon Ultra centrifugal filter units (Millipore). Amicon Ultra centrifugal filter units were equilibrated with 15ml of 0.2µm filtered DPBS at 3000xg at 4°C for 30 min. To concentrate supernatants suspension was loaded onto Amicon ultra centrifugal filter units and centrifuged at 3000rpm for 30min at 4°C. Each concentrated sample was collected at a final volume of 500µl. Human serum extracellular vesicles and vesicles from MEF cell supernatants were isolated using Izon original or Izon qev2 columns (IZON). The columns were equilibrated with 3 volumes of DPBS and 500µl sample was added on top of the columns. Void volumes of 3ml were let to pass, the subsequent 1.5ml fractions containing EVs were collected. The fractions were pooled and concentrated using 3kDa Amicon Ultra centrifugal units (Millipore) by centrifugation at 3000rpm for 30 min.

### Nanoparticle Tracking Analysis (NTA)

Isolated extracellular vesicle concentration was determined using Nanoparticle tracking analysis (NTA). The samples containing extracellular vesicles were diluted 1:500 or 1:500000 in 0.2µm filtered DPBS, then applied to the nanoparticle tracker (ZETA VIEW particle tracking analyzer) by syringe. Samples were measured in triplicates; the machine was set to 3 measurements at 11 positions for each replicate by the software Zeta View version8.05.12 SP1. Particle concentration was multiplied with dilution factor and normalized to serum volume used for EVs isolation.

### Extracellular vesicle DNA isolation and quantification

For isolation of DNA from inside extracellular vesicles, extracellular vesicle samples were depleted of extravesicular DNA using Benzonase nuclease (100 IU/ml), MgCl2 (2.5 mM) and CaCl2 (100 μM) at 37°C for 30min. Benzonase was inactivated using 1mM EDTA. Vesicle DNA was then isolated using AMPure XP beads for EV DNA isolation and purification (Beckman coulter) according to manufacturer instructions and resuspended in 100µl ddH_2_O. DNA concentrations were measured using nano-drop.

### Extracellular vesicle induced inflammation in mice and humans

*Sting^+/+^* and *Sting^-/-^* BMDMs were either stimulated with mouse plasma 1.35×10^11^ *Lrrk2^GoF^*-derived EVs or plasma PD-derived EVs corresponding to 15μl of human plasma (EVs were isolated from each patient separately) for 6h. After stimulation, the cells were washed with PBS and lysed for RNA isolation and preparation for qRT-PCR analyses. In an alternative protocol *Sting^+/+^* BMDMs where pre-treated with 15µM STING inhibitor (Invitrogen, H-151) to block STING activity before stimulation.

### Cellular senescence (SA-β-Gal assays)

To study cellular senescence the proliferation of WT and *Lrrk2^GoF^* MEFs were counted by trypan blue exclusion every third day and cells were stained for β-galactosidase activity with the CellEvent^TM^ Senescence Green Flow Cytometry Assay kit (Invitrogen. #C10840) according to manufacturer’s instructions.

### Purification of Cytoplasmic DNA

Cytoplasmic extract (CytExt) was isolated as previously described with minor modifications [76]. Briefly, MEFs were lysed in 10 mM HEPES (pH 7.9), 10 mM KCl, 1.5 mM MgCl2, 0.34 M sucrose, 10% (v/v) glycerol, plus protease inhibitors for 5 min on ice with 0.1% (v/v) Triton X-100, and nuclei were removed by low-speed centrifugation (1,500 × g, 10 min). Cytoplasmic protein extracts were treated with 1 mg/ml Proteinase K at 55°C for 1 hr. After phenol/chloroform extraction, the aqueous supernatant was incubated with 500 μg/ml DNase-free RNase A (QIAGEN) for 30 min at 37°C, again followed by phenol/chloroform extraction. The DNA-containing aqueous phase was precipitated, resuspended in TE buffer, DNA concentrations were adjusted to the protein concentration of cytoplasmic fractions and analyzed by a 2% (w/v) agarose gel in 1 × TBE buffer with 1 × GelRed stain (Biotium, Madison, WI) incorporated in gel.

### Real time qPCR analysis

Total RNA was isolated from tissues and cells using either EZNA HP RNA isolation kit (Omega, #R6812-02) or RNeasy Mini kit (Qiagen, #74104) or Trizol (Invitrogen, #15596026) according to manufacturer instructions. RNA was reverse transcribed using reverse transcription kit (Quantitect #205313 or Applied Biosystems #4368813). qRT-qPCR analysis was done on the 7500 real time PCR system (Applied biosystems) using Taqman Universal Master Mix (Thermo Fisher Scientific, #4440040) or Power Sybrgeen Master Mix (Thermo Fisher Scientific, #4367659) depending on primers used. The results were normalized to *Tbp1*/*TBP1* (reference gene) and expressed as fold change relative to untreated wild type control using CT values.

### Bulk RNA sequencing

RNA for sequencing was extracted from homogenized spleen, microglia and BMDMs using Opti-prep RNA isolation kit (Invitrogen, #AM1924). RNA was sent to Novogene (UK) for RNA-seq. Genes related to type I IFNs and cellular senescence were extracted and plotted as log2fold change compared to WT control.

### Phago/ Endolysosomal proteolysis assay

For phago/endolysosomal proteolytic assay, macrophages or MEFs were seeded into a 384-well plate and treated with 3.0µm or 0.3µm silica beads (Kisker Biotech) coated with DQ Green BSA (Thermo Fisher Scientific). Beads were diluted at 1:100 by binding buffer (1 mM CaCl_2_, 2.7 mM KCl, 0.5 mM MgCl_2_, 5 mM dextrose, 10 mM Hepes, and 5% FBS in PBS, pH 7.2) and kept with cells for 5min. After treatment the beads were replaced by warm binding buffer and real-time fluorescence was measured at 37°C on a SpectraMax i3x plate reader using excitation/emission wavelengths of 470/525 nm.

### Western blot analysis

For western blot cells were lysed using 200µl of 2x Laemmli buffer. Protein concentrations for western blot were determined using Bradford assay, 15µg of protein was used for immunoblotting. Proteins were separated on 4-12% SDS-PAGE gels and immunoblotted onto nitrocellulose or PVDF membranes (GE healthcare cat#10600023). Membranes were blocked in 5% blocking milk or 1x Roti Block (Roth) for 1h, then incubated in primary antibody (CD81, Histone H3) over night at 4°C. After incubation with horse radish peroxidase (HRP)-conjugated secondary antibody (α-Rabbit, α-Mouse) for 1h, proteins were detected using ECL substrate (Biorad) and visualized using photosensitive X-ray-film.

### Quantification and statistical analysis

Statistical analyses were done using GraphPad prism software versions 10. Data in text and figures are represented as mean ± SEM. Statistical comparisons were made using either one-way ANOVA followed by Tukeýs post hoc test, or students T-tests, P-values < 0.05 were considered significant.

## Acknowledgements

This work was funded by the University of Gothenburg and Knut and Alice Wallenberg Foundation to A.H., The Swedish Research Council (2023-01962 to AH, 2016-00890 and 2022-01308_3 to N.OG), The Swedish Cancer Foundation (22 2461 Pj 01H to A.H and CAN 20 1252PjF01H, CAN 23 3096 Pj, to N.O.G.) and STINT (CS2018-7952, to A.H.)

## Author contribution

A.H. conceived, designed, supervised the study. N.O.G conceived and designed experiments. M.Ö. performed most of the experiments. M.Ö. analyzed human PD patient and healthy donor data. M.Ö., I.M. performed behavioral studies, isolated mouse cerebrospinal fluid. M.Ö., A.S. performed flow cytometry data. M.Ö., C.M., N.S., D.F., I.F., L.R.G. performed in vitro experiments. K.S. provided tools for behavioral studies. M.A.K. provided PD patients and healthy donor-derived CSF extracellular vesicles. G.P. provided PD patients and healthy donor human patient plasma. M.Ö., N.O.G. and A.H. prepared the manuscript with comments from the other authors.

## Additional information

**Supplementary Information** is available for this paper.

**Correspondence and requests for** materials should be addressed to Anetta Härtlova.

