## Supplementary Figures for "Aging-associated endolysosomal decline drives inflammaging and neurodegeneration through the STING-IFN-I axis"

### Extended Data Figure 1

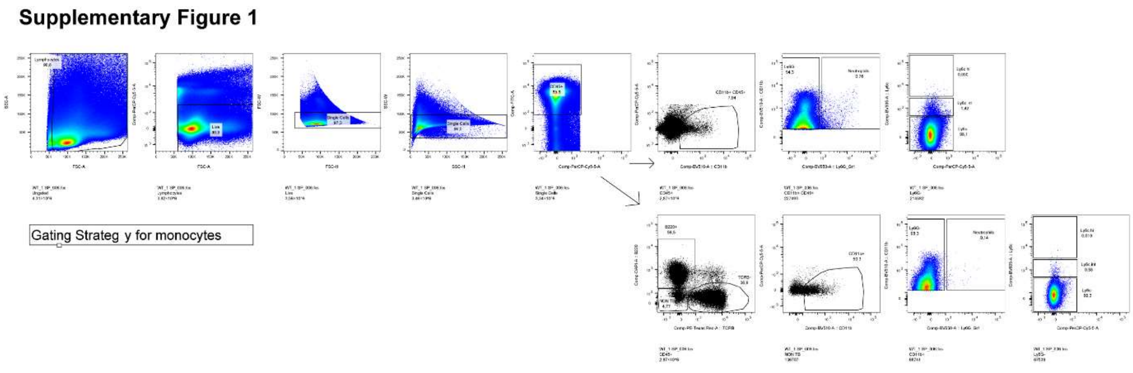


**Extended Data Fig. 1:** Flow cytometry strategy gating strategy for blood monocytes.

#### Extended Data Figure 2

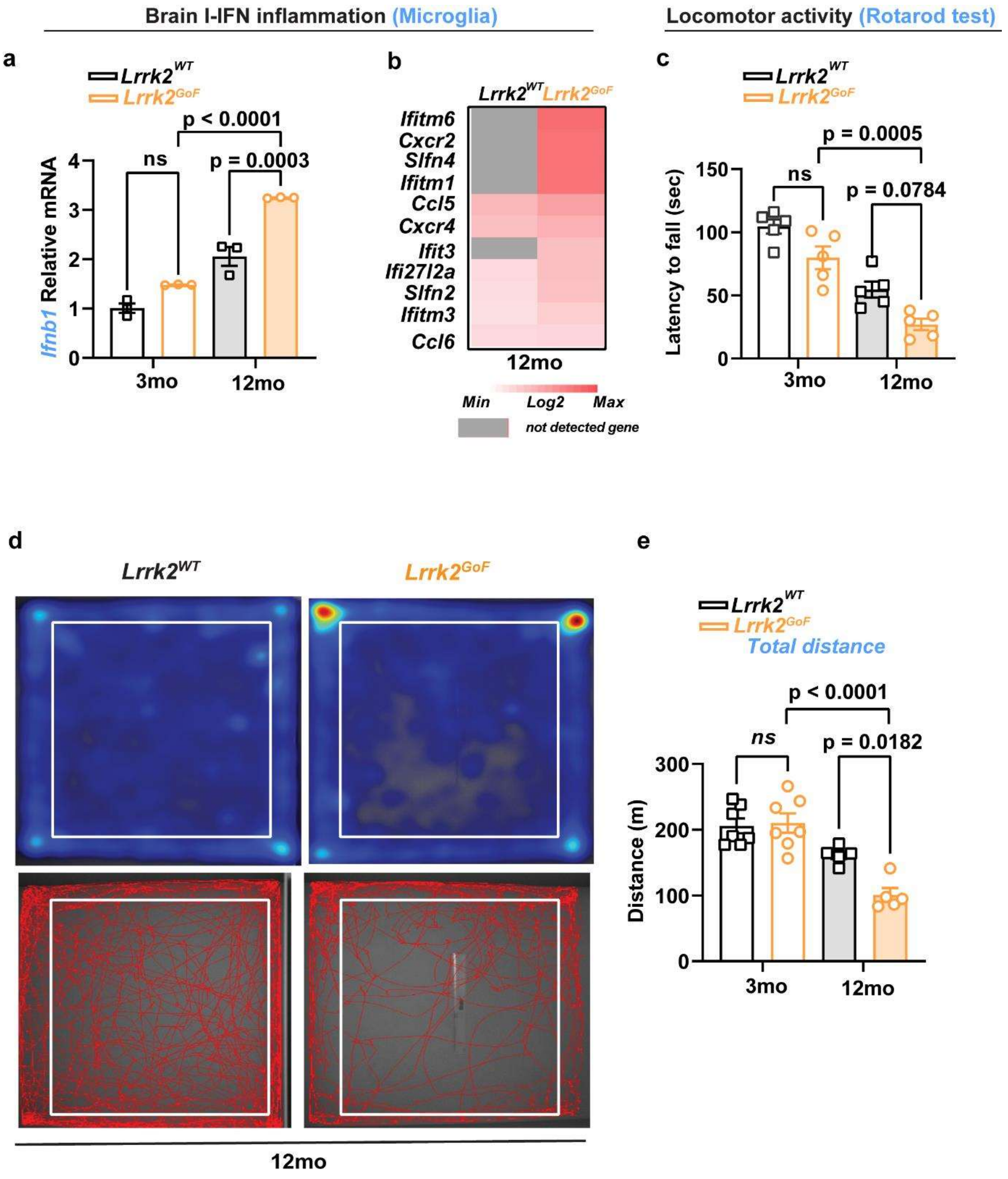

**Extended Data Fig. 2: *Lrrk2$^{GoF}$*mice exhibit accelerated age-associated brain inflammation and neurological decline. (A)** *Ifnb1* transcripts in microglia of 3 and 12 months old *Lrrk2$^{WT}$* and *Lrrk2$^{GoF}$*mice (n=3 mice per group). **(B)** Heatmap showing differently express type I-IFN/inflammatory in microglia of 3 and 12 months old *Lrrk2$^{WT}$* and *Lrrk2$^{GoF}$*mice (n=3 mice per group). **(C)** Rotarod test showing locomotor activity of 3 and 12 months old *Lrrk2$^{WT}$* and *Lrrk2$^{GoF}$* mice (n=5 mice per group). **(D)** Representative images of track plot reports recorded during the test sessions (ANY-maze) of aged *Lrrk2$^{WT}$* and *Lrrk2$^{GoF}$*mice. The central square was denominated "central zone" and the periphery, "border zone" (n=5 mice per group). **(E)** Locomotor activity of 3 and 12 months old *Lrrk2$^{WT}$* and *LRRK2$^{GoF}$* mice measured as total distance travelled within open field area (n=7 mice per young group, n=5 mice per aged group)**.** Statistics: one-way ANOVA followed by Tukey's post hoc test **(A, C, F)**. Error bars represent mean ± SEM.

**Extended Data Figure 3**

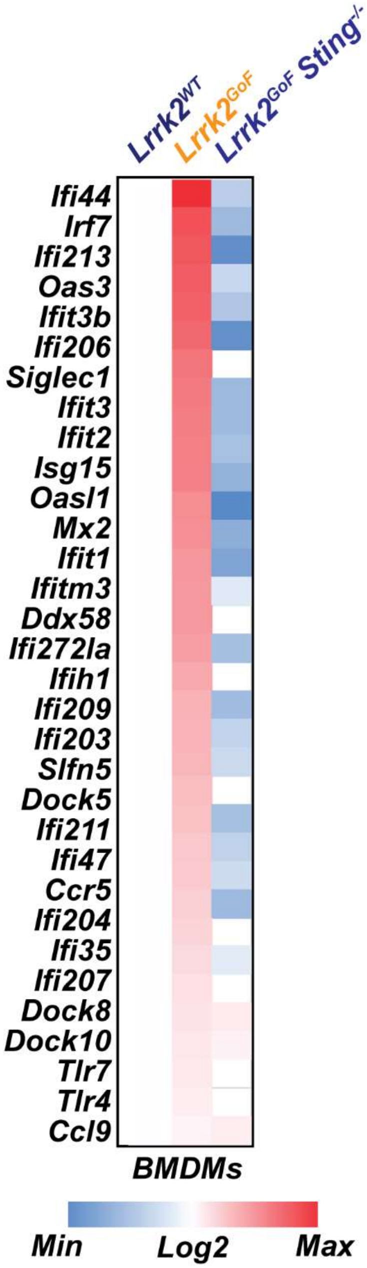


**Extended Data Fig. 3:** Heatmap comparing mean of differently expressed type I-IFN/inflammatory genes in *Lrrk2*$^{WT}$, *Lrrk2*$^{GoF}$ and *Lrrk2*$^{GoF}$*Sting*$^{-/-}$ BMDMs.

**Extended Data Figure 4**

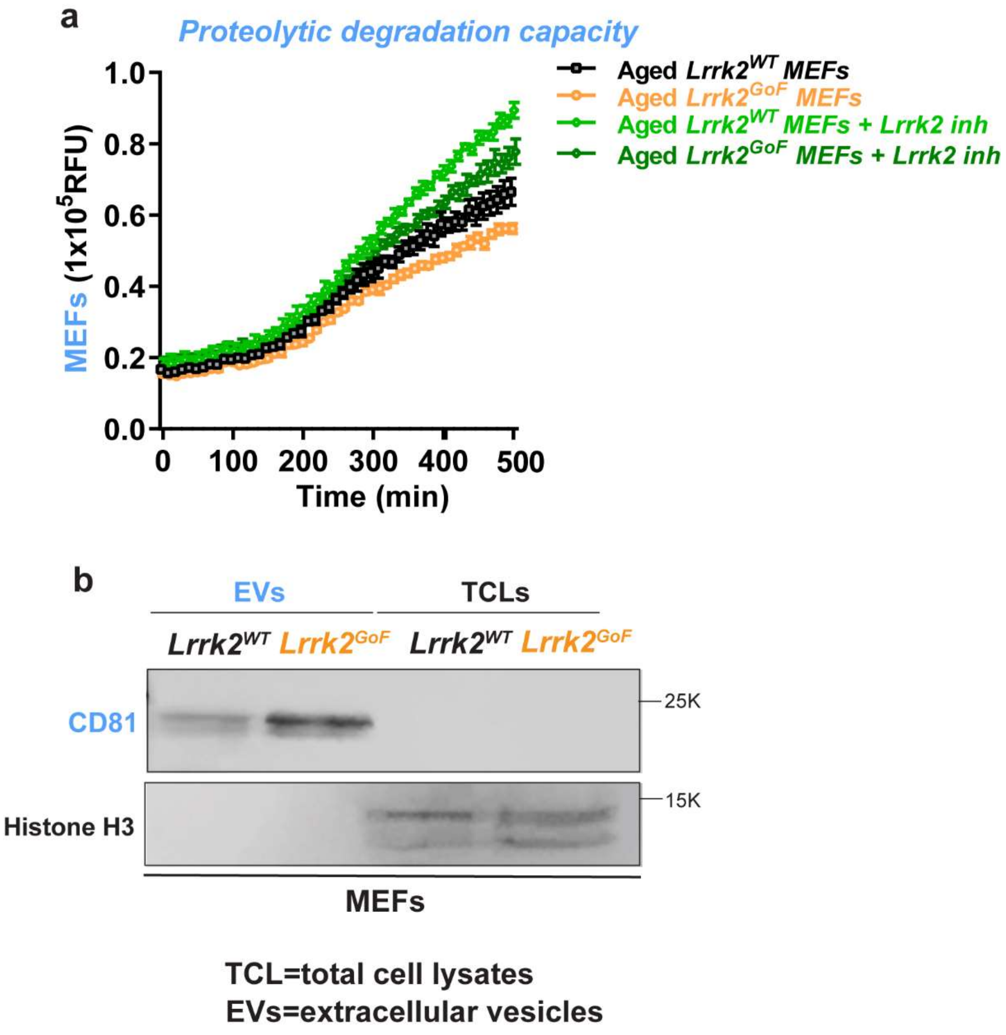


**Extended Data Fig. 3: (a)** Endo-lysosomal BSA DQ-green degradation in aged (senescent) WT and *Lrrk2^GoF* MEFs pre-treated or not with 100 nM Lrrk2 inhibitor MLI-2 (n=3 per group) **(b)** Western blot analysis of. extracellular vesicles (EVs) and total cell lysates isolated from WT and *Lrrk2^GoF* MEFs blotted for CD81 (Extracellular vesicle marker) and Histone H3 (nucleus).

**Extended Data Figure 5**

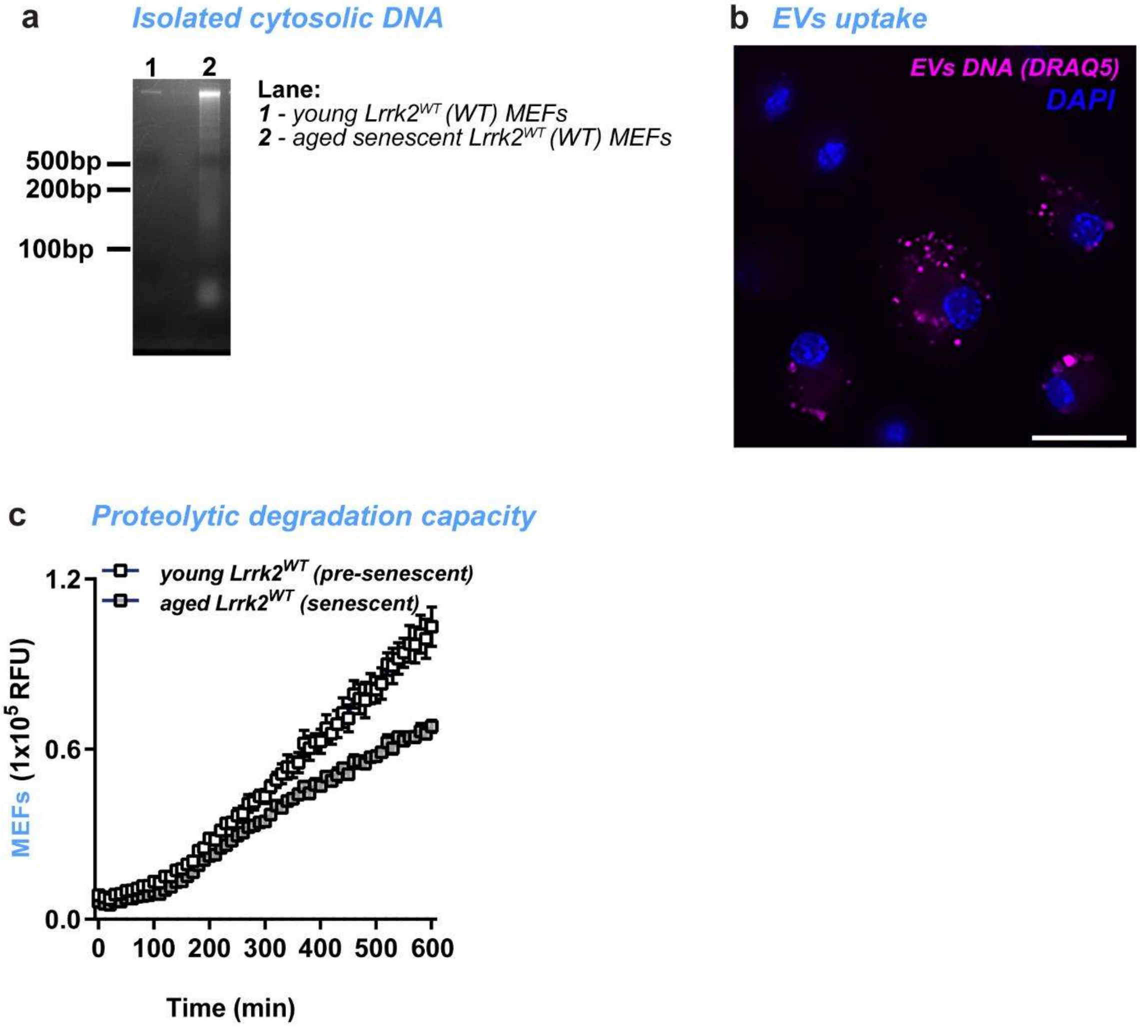


**Extended Data Fig. 5: (a)** Isolated cytoplasmic DNA of young (pre-senescent) and aged (senescent) WT MEFs. Error bars represent mean ± SEM. (**b**) Representative microscopy picture of intracellular PD-derived EVs containing DNA labeled by DRAQ5 dye in WT BMDMs represented by DAPI, Scale: 20μM. **(c)** Endo-lysosomal BSA DQ-green degradation in young (pre-senescent) and aged (senescent) WT MEFs. (n=3 per group).

#### Extended Data Figure 6

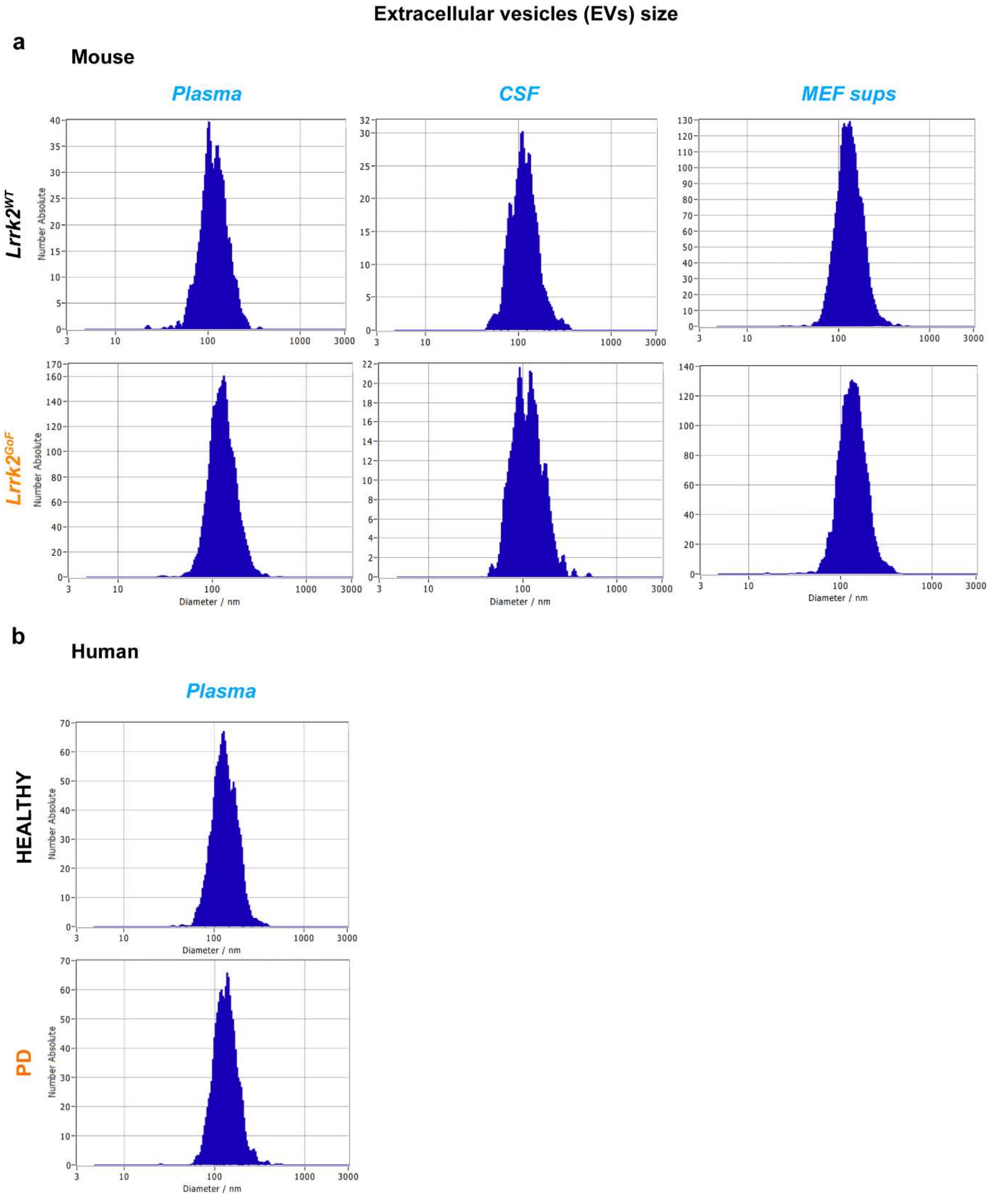


**Extended Data Fig. 6:** Size-characterization of enriched extracellular vesicles (EVs) from mouse plasma, csf and *in-vitro* MEFs cultures isolated from *Lrr2*$^{WT}$ and *Lrrk2* $^{GoF}$ mice (**a**) and

EVs enriched from human healthy donor and PD patient plasma and csf (**b**) by nanoparticle tracking analyzer (NTA).

#### Extended Data Figure 7

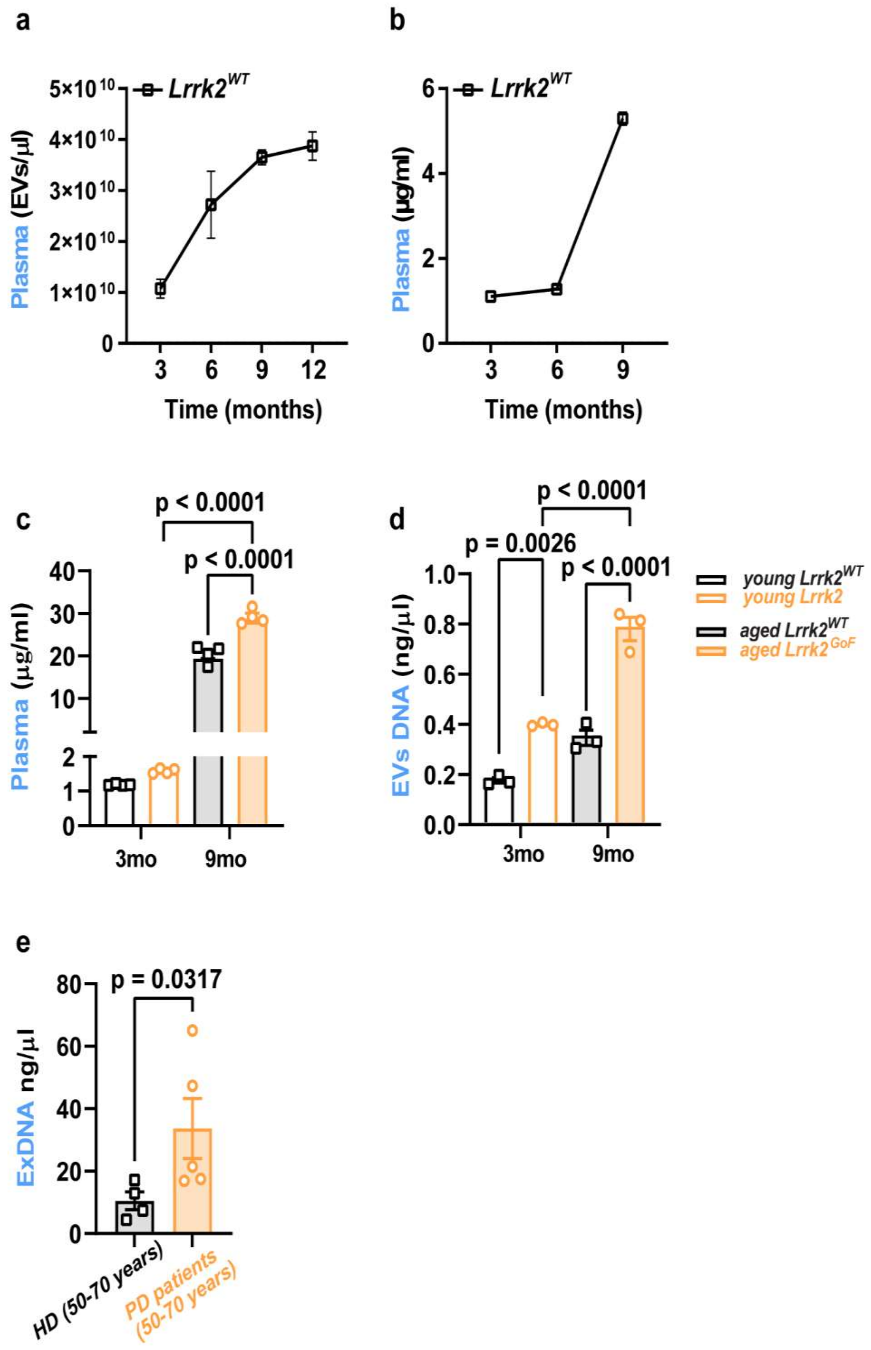


**Extended Data Fig. 7*:* (a-b)** NTA quantification and protein concentration of extracellular vesicles isolated from plasma of 3, 6, 9, 12 months old wild-type mice. (n=3 donors per group)

**(c)** Protein concentration of extracellular vesicles isolated from plasma of young and aged *Lrrk2*$^{WT}$ and *Lrrk2*$^{GoF}$*.* (n=3 donors per group) **(d)** Concentration of DNA isolated from the extracellular vesicles from plasma of young and aged *Lrrk2*$^{WT}$ and *Lrrk2*$^{GoF}$*.* (n=3 donors per group) **(e)** Concentration of DNA isolated from the extracellular vesicles from plasma of healthy donors and PD patients. (n=5 donors per group) Statistical tests: two-sided Student's t-test **(e)**, one-way ANOVA followed by Tukey's post hoc test **(c-d)**. Error bars represent mean ± SEM.
