## Supplementary material for "Aging-associated endolysosomal decline drives inflammaging and neurodegeneration through the STING-IFN-I axis": Resource Table

**Material and methods**

| REAGENT or RESOURCE | SOURCE | IDENTIFIER |
|---|---|---|
| **Antibodies** | | |
| CX3CR1-FITC | BioLegend | Cat#149019 |
| Clec12A-PE | BioLegend | Cat#143403 |
| CD11b-PE-cy7 | BD Biosiences | Cat#552850 |
| CD206-AF647 | BioLegend | Cat#141711 |
| CD45-BV425 | BioLegend | Cat#103133 |
| CD11c-BV711 | BioLegend | Cat#117349 |
| MHCII-A700 | Thermo Fischer Scientific, Invitrogen | Cat#56-5321-80 |
| CD86-PE | BD Biosienes | Cat#553692 |
| Ly6c-BV605 | BD Bioscienses | Cat#563011 |
| Anti-Rabbit HRP conjungated secondary ab | Cell Signaling Technology | Cat#7074S |
| Anti-Mouse HRP conjungated secondary ab | Cell Signaling Technology | Cat#7076 |
| Tyrosine Hydroxylase | Bioteche | Cat#NB300-109-25UL |
| Goat-anti-Rabbit Cy5 | Jackson Laboratory | Cat#111-175-003 |
| **Bacterial and virus strains** | | |
| Herpes simplex virus type I strain KOS KOS/Dlux/OriL (HSV-I-luc) containing a firefly luciferase reporter | The laboratory of Nelson O. Gekara, Stockholm University, Stockholm, Sweden | N/A |
| **Biological samples** | | |
| Human blood samples | The laboratory of Gesine Paul, Lund University, Lund, Sweden | N/A |
| Human CSF | Marzena Akanbi-Kurzawa, Biosciences Institute, Newcastle University, Newcastle upon Tyne, UK | N/A |
| **Chemicals, peptides, and recombinant proteins** | | |
| Dulbecco's Modified Eagle Medium (DMEM) | Thermo Fisher Scientific, Gibco | Cat#31966-021 |
| Iscove's Modified Dulbecco's Medium (IMDM) | Thermo Fisher Scientific, Gibco | Cat#12200-069 |
| Fetal bovine serum (FBS) (heat-inactivated) | Thermo Fisher Scientific, Gibco | Cat#26140-087 |
| Extracellular vesicles (exosomes) depleted Fetal bovine serum (FBS) | Thermo Fischer Scientific, Gibco | Cat#A2720803 |
| Penicillin-streptomycin | Thermo Fisher Scientific, Gibco | Cat#15140-122 |
| Benzonase nuclease | EMD Millipore Company | Cat#71206-25KUN |
| DNase I | Roche | Cat#11 284 932 001 |

| | | |
|---|---|---|
| TaqMan Gene Expression Master Mix | Thermo Fisher Scientific | Cat#4304437 |
| Power SYBRgreen PCR master mix | Thermo Fisher Scientific | Cat#43067659 |
| RNeasy Mini kit | Qiagen | Cat#74104 |
| RNA isolation kit | Omega | Cat#R6812-02 |
| RNA isolation kit -optiprep | Invitrogen | Cat#AM1924 |
| Adult Brain dissociation kit | Miltenyi Biotec | Cat#130-107-677 |
| CD11b positive microbeads human | Miltenyi Biotec | Cat#130-049-601 |
| CD14 positive microbeads human | Miltenyi Biotec | Cat#130-050-201 |
| CD11b positive microbeads mouse | Miltenyi Biotec | Cat#130-126-725 |
| Amicon Ultra-2 Centrifugal Filter Units | Merck Millipore, Sigma-Aldrich | Cat#UFC200324 |
| Amicon Ultra-15 Centrifugal Filters (30kDa MWCO) | Merck Millipore, Sigma-Aldrich | Cat#UFC903024 |
| Amicon Ultra-15 Centrifugal Filters (10kDa MWCO) | Merck Millipore, Sigma-Aldrich | Cat#UFC901024 |
| Pierce BCA protein assay | Merck Millipore, Sigma-Aldrich | Cat#QPBCA |
| Bradford coomassie plus protein assay | Thermo Fischer scientific | Cat#1856210 |
| Ponceau S solution | Biotium | Cat#22001 |
| Collagenase IV | Merck Millipore, Sigma-Aldrich | Cat#C4-BIOC |
| Tween20 | Merck Millipore, Sigma-Aldrich | Cat#SLCC6187 |
| DMSO | Merck Millipore, Sigma-Aldrich | Cat#8590 |
| Recombinant mouse IFNb | R&D Systems | Cat#8234-MB |
| Recombinant human IFNb | R&D Systems | Cat#8499-IF |
| DAPI | Merck Millipore, Sigma-Aldrich | Cat#28718-90-3 |
| Hoescht | Thermo Fisher scientific | Cat#33342 |
| Zeocin | Invivogen | Cat#ant-zn |
| Normocin | Invivogen | Cat#ant-nr |
| TRIzol Reagent | Thermo Fischer Scientific, Invitrogen | Cat#15596026 |
| Crystal Violet | Merck Millipore, Sigma-Aldrich | Cat#HT90132-1L |
| Brilliant Stain Buffer | BD Horizon | Cat#563794 |
| Roti block | Roth | Cat#A151.1 |
| DRAQ5 | Thermo Fischer Scientific | Cat#62254 |
| Fluorescence mounting medium | Agilent Dako | Cat#S3023 |
| Triton-X-100 | Merck Millipore, Sigma-Aldrich | Cat#1002538786 |
| RNaseA | Qiagen | Cat#19101 |
| Proteinase K | Qiagen | Cat#RP103B- |
| TE buffer, 1xsolution | Thermo Fischer Scientific | Cat#J75893.XCR |
| **Critical commercial assays** | | |

| | | |
|---|---|---|
| qEV2 / 70 nm columns for isolation of membrane vesicles | IZON | Cat#IC2-70 |
| AMPure XP beads for EV DNA isolation and purification | Beckman Coulter | Cat#A63881 |
| QuantiTect Reverse Transcription Kit | Qiagen | Cat#205311 |
| cDNA synthesis kit | Thermo Fischer Scientific, Applied Biosystems | Cat#4368813 |
| CellEvent™ Senescence Green Flow Cytometry Assay Kit | Thermo Fischer Scientific, Invitrogen | Cat#C10840 |
| Senescence β-Galactosidase Staining Kit | Cell Signaling Technology | Cat#9860 |
| Serum/plasma extracellular vesicles (exosomes) isolation kit | Qiagen | Cat#76603 |
| Cell medium/ urine / csf extracellular vesicles (exosomes) isolation kit | Qiagen | Cat#76743 |
| **Experimental models: Cell lines** | | |
| L929 (NCTC clone 929) | ATCC | Cat#ATCC:CCL-1 |
| WT (C57BL/6NTac) and Lrrk2$^{GoF}$ (C57BL/6-Lrrk2tm4.1Arte) mouse embryonic fibroblasts | This Manuscript | |
| B16 Blue ISG | InvivoGen | Cat#bb-ifnabg |
| HEK-Blue™ ISG cells | InvivoGen | Cat#hbg-isg-1 |
| **Experimental models: Organisms/strains** | | |
| WT (C57BL/6NTac) mice | Taconic | Model#B6 |
| *Lrrk2*$^{GoF}$ (G2019S KI) (C57BL/6-Lrrk2tm4.1Arte) mice | Taconic | Model#13940 |
| *Sting*$^{-/-}$ (C57BL/6J-Tmem173gt/J) | The Jackson Laboratory | Strain#017537 RRID: IMSR_JAX:017537 |
| *Ticam*$^{-/-}$(C57BL/6J-Ticam1Lps2/J) | The Jackson Laboratory | #005037 |
| *Lrrk2*$^{GoF}$ *Sting*$^{-/-}$ | This Manuscript | |
| *Lrrk2*$^{GoF}$ *Ticam*$^{-/-}$ | This Manuscript | |
| **Oligonucleotides** | | |
| **Mouse Tbp1 primer1:** **Fw:** 5'-GAAGCTGCGGTACAATTCCAG-3' | Harvard primer bank | Invitrogen |
| **Mouse Tbp1 primer2:** **Rv:** 5'-CCCCTTGTACCCTTCACCAAT-3' | Harvard primer bank | Invitrogen |
| **Mouse TaqMan™ Gene Expression Assay (FAM)** (*Ifnb1*), Assay ID: Mm00439552_s1 | Thermo Fischer Scientific, Applied Biosystems | Cat#4331182 |
| **Mouse Quantitect Primer Assay Tnfa,** Mm_TNF_1_SG QuantiTect Primer Assay | Qiagen | Cat#QT00104006 |
| **Human TBP1 primer1:Fw:** 5'-GATAAGAGAGCCACGAACCAC-3' | Harvard primer bank | Invitrogen |
| **Human TBP1 primer2:** **Rev:** 5'-CAAGAACTTAGCTGGAAAACCC-3' | Harvard primer bank | Invitrogen |

| | | |
|---|---|---|
| **Human Quantitect Primer Assay TNFA,** Hs_TNF_1_SG QuantiTect Primer Assay | Qiagen | Cat#QT00029162 |
| **Human Quantitect Primer Assay IFNB1** Hs_IFNB1_1_SG QuantiTect Primer Assay | Qiagen | Cat#QT00203763 |
| **Software and algorithms** | | |
| GraphPad Prism softwere version 9.5.0 and 10 | GraphPad | https://www.graphpad.com/scientific-software/prism/ |
| FlowJo software version 10.8.1. | FlowJo | https://www.flowjo.com/ |
| Zetaview software version 8.05.10 SP1 | Particle Metrix | https://particle-metrix.com/zetaview/ |
