## Supplementary Table 1 for "Aging-associated endolysosomal decline drives inflammaging and neurodegeneration through the STING-IFN-I axis"

| **Sample-Id** | **Gender** | **Age** | **Health status** | **Mutation** |
|---|---|---|---|---|
| **1** | Female | 30 | HD | |
| **2** | Female | 26 | HD | |
| **3** | Female | 25 | HD | |
| **4** | Male | 30 | HD | |
| **5** | Female | 28 | HD | |
| **6** | Male | 53 | HD | |
| **7** | Male | 66 | HD | |
| **8** | Male | 67 | HD | |
| **9** | Female | 56 | HD | |
| **10** | Female | 54 | HD | |
| **11** | Female | 53 | PD | LRKK2 G2019S |
| **12** | Female | 65 | PD | |
| **13** | Female | 58 | PD | |
| **14** | Male | 59 | PD | |
| **15** | Male | 53 | HD | |
| **16** | Female | 52 | HD | |
| **17** | Female | 24 | PD | Parkin hom |

**Supplemental Table 1*:*** Overview of age, gender of the Parkinson´s patients and healthy donors.
